# Measuring the double-drift illusion and its resets with hand trajectories

**DOI:** 10.1101/2021.08.06.455415

**Authors:** Bernard Marius ’t Hart, Denise Y.P. Henriques, Patrick Cavanagh

**Affiliations:** Centre for Vision Research, York University, Toronto, ON M3J1P3, Canada; Department of Psychology, Glendon College, Toronto, ON M4N3M6, Canada

## Abstract

If a gabor pattern drifts in one direction while its internal texture drifts in the orthogonal direction, its perceived position deviates further and further away from its true path. We first evaluated the illusion using manual tracking. Participants followed the gabor with a stylus on a drawing tablet that coincided optically with the horizontal monitor surface. Their hand and the stylus were not visible during the tracking. The magnitude of the tracking illusion corresponded closely to previous perceptual and pointing measures indicating that manual tracking is a valid measure for the illusion. This allowed us to use it in a second experiment to capture the behavior of the illusion as it eventually degrades and breaks down in single trials. Specifically, the deviation of the gabor stops accumulating at some point and either stays at a fixed offset or resets toward the veridical position. To report the perceived trajectory of the gabor, participants drew it after the gabor was removed from the monitor. Resets were detected and analyzed and their distribution matches neither a temporal nor a spatial limit, but rather a broad gamma distribution over time. This suggests that resets are triggered randomly, about once per 1.3 *s*, possible by extraneous distractions or eye movements.

## Introduction

Encoding the position of objects in the world is necessary for the myriad of visually guided motor tasks that we do every day. It is no surprise then that the brain has several mechanisms to gauge and update the position of objects in the world. For example, when a target is moving, its motion can influence its perceived location, making it appear to be slightly ahead of its true location, e.g. the flash lag (Nijhawan, 1994); or the flash grab effect (Cavanagh & Anstis, 2013). Although there are other explanations for the flash-lag effect (Cai & Schlag, 2001; Eagleman & Sejnowski, 2000), these cases of position extrapolation may be functional, helping the motor system overcome inevitable neural delays in targeting a moving object (Duhamel et al., 1992; Hogendoorn, 2020; Nijhawan, 1994). A stronger and very different motion-induced position shift arises when a moving gabor has internal motion orthogonal to its path and is viewed in the periphery (Gurnsey & Biard, 2012; Kwon et al., 2015; Lisi & Cavanagh, 2015; Shapiro et al., 2010; Tse & Hsieh, 2006). This double-drift stimulus (Fig. 1, left) generates extreme misjudgments of the moving target’s location which may deviate by as much as several degrees of visual angle from its true location. Surprisingly, this extraordinary perceptual illusion does not affect eye movements to the gabor: immediate saccades to these targets are determined by their physical, not their perceived location (Lisi & Cavanagh, 2015). In contrast, delayed eye movements and pointing go to the perceived position (Lisi & Cavanagh, 2017; Massendari et al., 2018; Ueda et al., 2018). These findings suggest that both the true and illusory positions are available in the brain.

**Figure 1:**
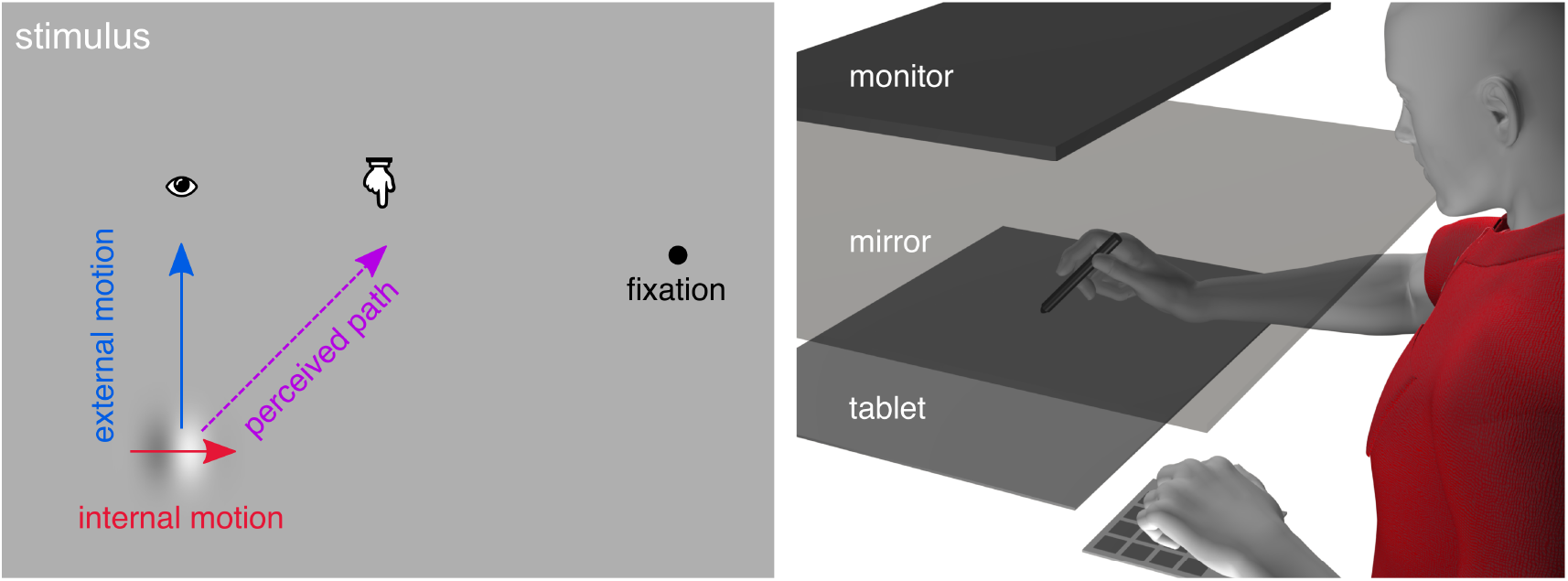
Stimuli and Setup **Left:** a gabor with internal motion (phase shifts) and external motion, that is viewed peripherally, appears to follow a path deviating from the true path. **Right:** participants track the perceived trajectory as the gabor moves back and forth along its path.

The strength of the double-drift illusion probably arises because of the very poor location information for a gabor pattern in the periphery when it has the same mean luminance as the background (Cavanagh & Tse, 2019; Gurnsey & Biard, 2012; Kwon et al., 2015). With poor positional certainty, the motion information contributes to the location estimate, generating a shift in perceived position. If the gabor itself is not moving (De Valois & De Valois, 1991), the shift saturates quickly (40 *ms*: Chung et al., 2007; 100 *ms*: Arnold et al., 2007), perhaps because the accumulating evidence of actual location of the stationary gabor patch is sufficient to anchor its position. However, when the gabor itself also moves, as in the double-drift stimulus, there is no stable location information to anchor the position estimate and the build-up continues well beyond 100 ms. The perceived location may move from the physical position by up to several times the size of the gabor.

At some point, the illusion does stop accumulating. The accumulation can be reset by introducing a temporal break (Lisi & Cavanagh, 2015) or by distracting attention (Nakayama & Holcombe, 2020), but it may also reset spontaneously, perhaps when the accumulation has gone on too long or too far. Informally, some observers have reported that the gabor’s path may saturate so that it remains parallel to, but offset from the true path. Others report that the position moves back toward its physical location, either slowly or suddenly, whereupon accumulation begins again. The point where the accumulation of illusory offset stops could be informative about how the brain combines motion and position information.

Here we first investigate wether hand-tracking of the gabor demonstrates the illusion. Eye and hand movement are differentially sensitive to many illusions, such as the Ponzo illusion (Gamble & Song, 2017), Ebbinghaus illusion (Knol et al., 2017), flash-lag effect (Blohm et al., 2003), the Duncker illusion (Soechting et al., 2001) and many others. Similarly, it has been shown that the double-drift illusion affects pointing movements, but not saccades (Lisi & Cavanagh, 2017), and hand movements have been used to test other perceptual phenomena (Patricio Décima et al., 2022). Hence, we expect that hand trajectories will be an effective method to capture the full illusory trajectory, rather than a point estimate of its endpoint (Hui et al., 2020) or its overall angle (Lisi & Cavanagh, 2015).

The strength of the illusory percept — the deviation of the perceived path of the gabor from its true path — depends on the speed of both the internal and external motion (Cavanagh & Tse, 2019; Heller et al., 2021; Kwon et al., 2015; Shapiro et al., 2010; Tse & Hsieh, 2006). In this first experiment, we test how well manual tracking reflects the percept of the illusion. To do so, we use hand movement trajectories recorded during stimulus presentation (Fig. 1). We compare trajectories across 5 different internal motions of the gabor and compare the illusion strength with previous work based on a straightforward vector combination model (Heller et al., 2021). As we expected, online tracking provides a measure of illusion strength that is similar to that found with pointing (Lisi & Cavanagh, 2017). However, the traces reveal only a few possible resets (>4% trials), and no limit to the illusion is clear.

To study resets, we therefore slow down the external stimulus motion in our second experiment, and have participants reproduce the perceived trajectory after stimulus presentation. Now we do observe spontaneous resets (64% of trials) that allow us to investigate the spatial and temporal limits of the illusion. We find that there is no hard temporal or spatial limit leading to resets. Instead, resets appear to occur randomly over time, based on a gamma distribution.

## Experiment 1: Online tracking

In this experiment, participants were asked to use a stylus on a drawing tablet to track a moving gabor with some amount of internal motion that elicited the double-drift illusion. A variant on online tracking has been used before (Cormack, 2019) with a continuous task where the stimulus depended on the response. Here, we test if tracking the illusion continuously is a good measure of illusion strength, or if it prevents or decreases the illusion. Additionally, we use a relatively long trajectory to allow the double-drift illusion to exceed any perceptual limits it may have, so that we can observe what these are.

### Methods

#### Participants

For this experiment, 4 participants were recruited from the lab (3 female; ages 21 – 31, mean: 25.75) and they were naive to the purpose of the study. All participants reported being right handed and had either normal or corrected to normal vision. Procedures were in accordance with the Declaration of Helsinki (2003) and were approved by York’s Human Participants Review Committee. All participants provided prior, written, informed consent.

#### Setup

Participants used a stylus on a horizontal drawing tablet (Wacom Intuos Pro) to indicate where they perceived the location of a gabor (see Fig. 1, right), as well as a keypad for additional responses. Stimuli were displayed on a downward facing LCD monitor (30 Hz, 20”, 1680×1050, Dell E2009Wt), parallel with the drawing tablet. The stimuli were observed via an upward facing mirror placed exactly between the tablet and monitor, so that the stimuli appear to be in the same horizontal plane as the hand, and the tip of the stylus. Experiments were run in Python 2.7 with PsychoPy (Peirce et al., 2019).

For reporting degrees of visual angle that stimuli spanned (*dva*), we assumed the participants to be close to the setup, with their eyes at the height of the monitor. While there was no requirement to use the forehead rest, this is a reasonable assumption, since stimuli are only visible when relatively close to the setup. Nevertheless, reported *dva’s* are estimates of the size of stimuli.

#### Stimuli

The stimuli were phase-changing gabor patches with a sigma (*σ* or sd) of 0.43 *cm* or ∼0.59 *dva*, spatial frequency of 0.58 *c/cm* or ∼0.42 *c/dva*, vertical orientation, and Michelson contrast of 98.7%. The background luminance was 18.46 *c d* / *m*^2^ and the mean luminance of the gabor deviated from the background by less than 1%. By simultaneously moving the gabor envelope and drifting its internal sine wave, it becomes a ‘double-drift stimulus’ where the perceived path deviates from the real path. The gabor changed phase at different rates, called ‘internal motion’ here, specified in cycles per second (*cps*). The real displacements of the stimulus are along the Y axis of the monitor, keeping the X coordinate constant at the middle of the screen, so that a stimulus without internal motion would appear to move toward and away from the participant on the horizontal display.

The motion of the envelope was restricted to the central 13.5 *cm* (∼14.0 *dva*) and was set so that the stimulus would move from one end to the other of its path (a motion we call a ‘pass’) in 2 *s*, corresponding to a speed of ∼6.75 *cm/s* (or roughly 7.0 *dva/s*).

#### Procedure

In this experiment, participants used their unseen hand and stylus to do continuous online tracking of the double-drift stimulus while that moved back and forth along the Y-axis of the horizontal screen for 12 *s*. Before trial onset, the participant moved the stylus to the middle of the screen, and were given 1.5 *s* to fixate a point on 16.5 *cm* (∼22.5 *dva*) to the left or right of the centre of the screen. The gabor would then appear at the middle of the screen and start moving. Internal motion could be 3, 1, 0, -1 or -3 *cps* (each repeated 8 times). Since each moving gabor was shown for 12 *s*, and it moved from one end to another in *2* s, it completed 5 full and 2 half passes of the path. The internal motion of the gabor was inverted at the far and close ends of the physical path, where external motion was also inverted.

#### Analyses

Because there is no set reference point to gauge movement direction against after each direction change, we use instantaneous heading along the trajectory instead. We first segmented the tracking trajectories according to full stimulus passes (in between direction changes of the gabor), and removed the noisy tracking during the first and last half second, leaving 1 *s* trajectory segments. We then calculated the instantaneous heading (direction, disregarding velocity or distance) between all 32 raw samples for 31 heading samples per segment. This heading measure should depend on the illusion strength.

To determine if hand tracking produces an illusion similar to that seen in other studies, we first compare our results to those reported by Heller et al. (2021) who used similar displays. We also compare the illusory direction of the heading against the predictions of a simple vector combination model (Cavanagh & Tse, 2019; Heller et al., 2021; Tse & Hsieh, 2006). There are alternative models that could be considered, one from Kwon et al. (2015) and another from Johnston and Scarfe (2013). Kwon et al.’s (2015) Kalman filter tracking model assumes a saturation of the position shifts by 200 ms, something we do not observe here. Johnson and Scarfe’s (2013) model used a harmonic average to combine vectors, a process that gives more weight to vectors with lower magnitudes. This model was used to estimate the perceived direction of an array of stationary Gabors with various internal motions but is less suited to the double-drift stimulus that has only two vectors, one of which is a second-order motion (the motion of the Gabor itself). In the simple vector combination model we use here, the perceived direction is a vector combination of the external (*V*_*e*_) and internal (*V*_*i*_) motions weighted by a constant *K*. That is, the deviation from the physical direction is given by:

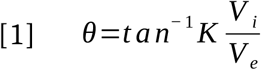

Here we calculate *V*_*i*_ and *V*_*e*_ in *cm/s*; *cps* * 0.58 *cm* for *V*_*i*_ and 13.5/2 *cm/s* for *V*_*e*_. If *K* is 0, then there is no illusion and the perceived direction matches the external motion direction. However, if *K* is 1, then the internal motion has equal contribution with the external motion in determining the perceived direction. For comparison with other work, we will find the value of *K* that best describes the strength of the illusion in each of our two experiments.

In particular, we compare the illusion strengths we find with those reported by Heller et al. (2021). They show a double-drift stimulus with 36 combinations of internal and external motion for 500 ms and then have participants indicate on a ring where they perceived the path of the stimulus to intersect with this ring. While they hypothesize that illusion strength decreases with faster external motion (perhaps because of resets or limits on perception), here we will use their overall estimate of illusion strength, given by *K* =0.74. While in *dva/s* our stimuli have relatively high external and relatively low internal motion, compared to Heller et al. (2021), they are within the range of tested stimuli as are the predicted illusion strengths.

All analyses were done in R 3.6.1 (R Core Team, 2019). All data, as well as scripts for the experiments and analyses are available on OSF: https://osf.io/72ndu/.

#### Results

All raw trajectories are shown in Figure 2A. In Figure 2C, we plot the average instantaneous heading during the middle second of each 2 *s* pass for each of the 5 internal motion speeds. The average angle of the tracked path is given in degrees deviation from straight forward. It seems that continuous tracking is sensitive to the strength of the illusion, and this is confirmed by a repeated-measures ANOVA on the average angles, using internal motion as a within-subjects factor (*F*(4,12)=102.6, *p*<.001, *η*^2^=0.97). With 4 FDR-corrected (Benjamini & Hochberg, 1995), paired t-tests, we find a difference in heading between internal motion speeds of 0 *cps* and both 1 *cps* (*p*=.022) and -1 *cps* (*p*=.025) as well as between -3 and -1 *cps* and 3 and 1 *cps* (both *p* <.001). This makes it clear that continuous tracking does not prevent the illusion from occurring. We will use heading angle as a measure of the strength of the illusion.

**Figure 2:**
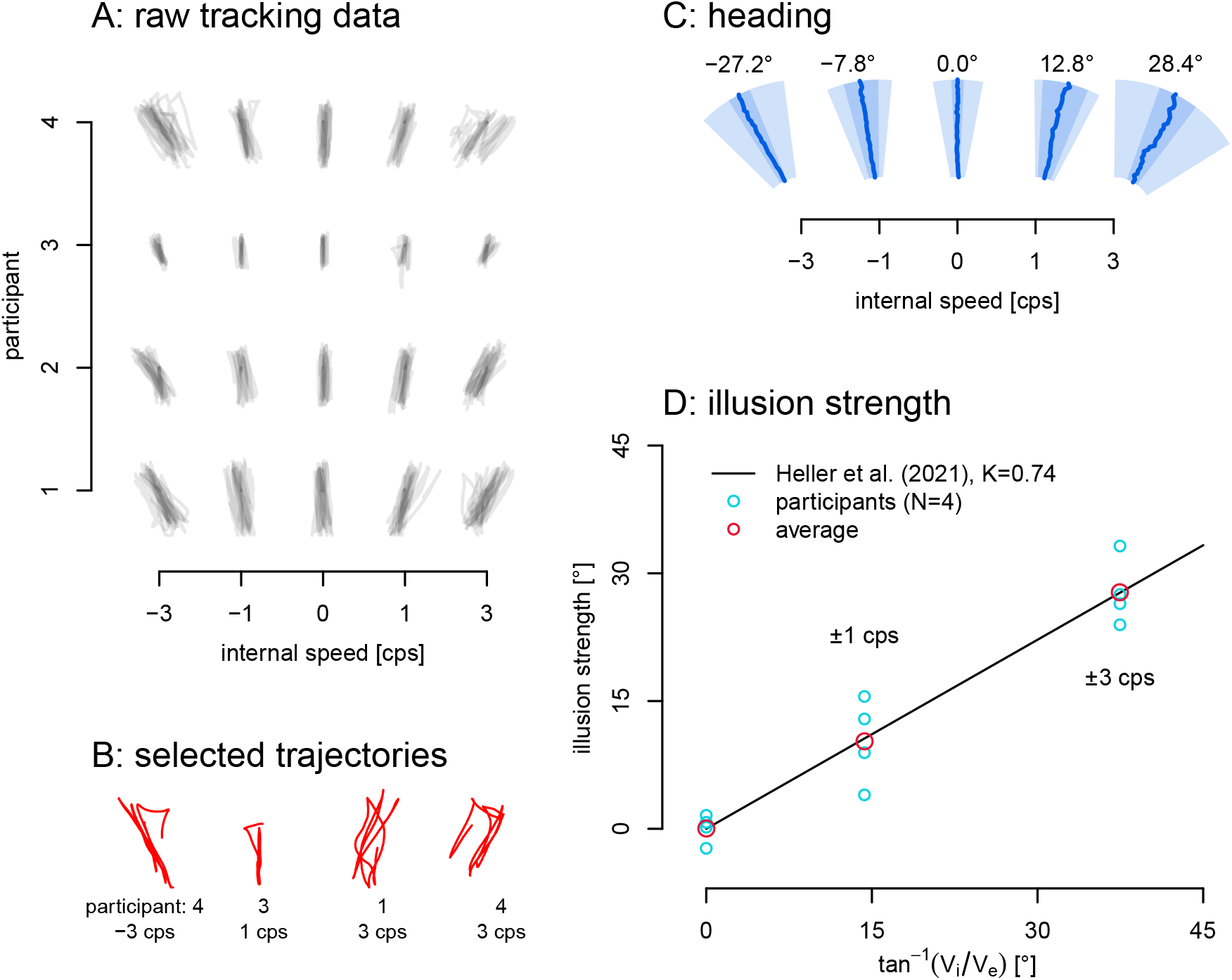
Online Tracking Data. **A:** Raw individual trajectories. In general, the traced paths approximate linearity, with the direction depending on the internal motion. **B:** Four sample trials with anomalous deviations that may be resets. **C:** Average heading of the middle 1 s of each pass. Distance from origin is time, angle is the heading angle. Dark blue lines show the average heading over time. The light blue depicts the average area (across participants) containing 90% of the heading data. The darker blue areas depict the 95% confidence interval of the mean heading. **D:** The strength of the illusion in degrees angle deviation from the physical path. In red are the average illusion strengths for 0 cps, ±1 cps and ±3 cps, and in light blue the data for the 4 individual participants. Black line: prediction of the vector combination model of Eq. 1 with K=0.74 as in Heller et al. (2021), which is indistinguishable from the best fit to our data: K=0.738.

#### Illusion strength

The illusion size we find with tracking appears to be in the range of that reported in other article for similar speeds and eccentricity. For example, Heller et al. (2021) reported an illusion of 30° for internal and external speeds of 7.2 and 7.2 dva/s compared to the 27.8° here (average of left and right 3Hz internal motion) with internal and external speeds both 7 dva/s. The data clearly show the illusion, and we can compare the mean illusion strengths here to those of Heller et al. (2021) in more detail using their vector combination model. The averages from our data here (Fig 2D: red circles) do coincide fairly well with the illusion strength predicted by the model fitted to their data. The best fit value of the vector combination model (Eq. 1 above; Fig 2D: black line) to our data *K* =0.738 is very close to *K*=0.74 reported by Heller et al. (2021). We can conclude that tracking the illusion while observing it does not appear to change its strength.

## Experiment 2: Delayed retracing

The illusion cannot drift continuously away from the physical path forever. It can be disrupted by a temporal break (Lisi & Cavanagh, 2015) or by distracting attention (Nakayama & Holcombe, 2020). These disruptions may cause the perceived path to stay at a fixed offset, travelling parallel to the physical path, or it may return toward the veridical position before resuming the illusory direction (Fig. 3). Other percepts of resets have been reported as well, but as far as we know these fall in between these two extremes. Such resets may occur spontaneously once the accumulation has gone on too long or too far, depending on what mechanism causes the resets, and the strength of the illusion (purple arrows in Fig. 3). The illusion strength in turn depends on the stimulus properties (speeds, gabor size, carrier frequency, eccentricity; Gurnsey & Biard (2012)) as well as variations within and between participants. The purpose of this experiment is to capture this spontaneous resetting and determine its source.

**Figure 3:**
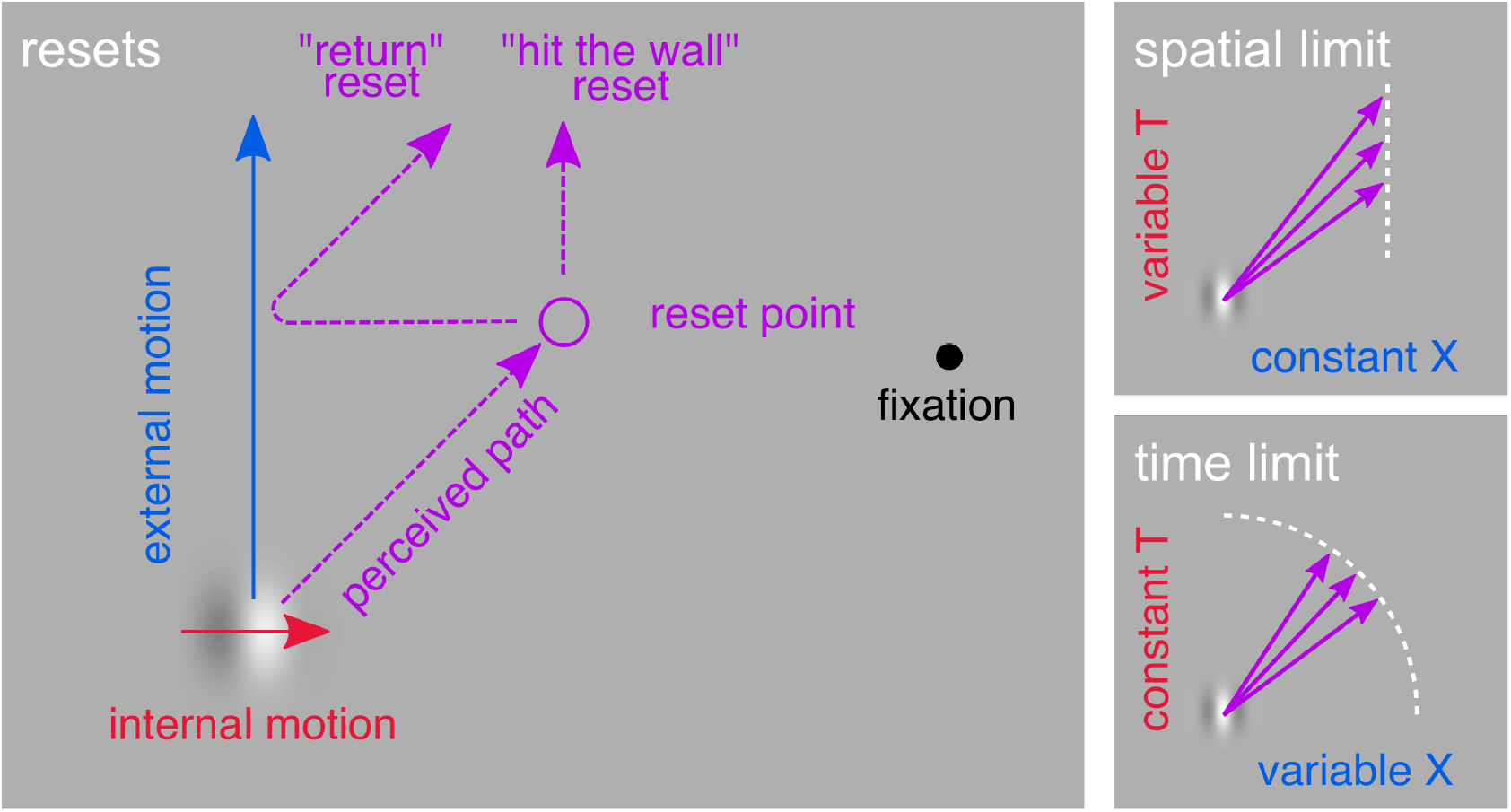
Double-Drift Resets. **Left:** At some point, the perceived position may stop moving further away from its actual path. For some people this takes the form of a ‘return’ reset, back toward the true position either suddenly or slowly, for others it means the illusory position remains at a fixed offset moving parallel to the real path. **Right:** Given varying strength of the illusion (denoted by the angle of the purple arrows), e.g. due to stimulus properties or within or between subject differences, resets may occur at different points in space. **Top right:** if these resets can be explained by a spatial limit on the size of the illusory position shift away from the true vertical path, the locations of the resets should have a constant X coordinate; the spatial offset from the vertical path. **Bottom right:** if they occur instead after some period of time, resets should have a constant radius (T); the time since the gabor’s motion began.

In the previous experiment with online tracking, the duration of each pass in the online tracking experiment was 2 seconds and the manual traces (Fig. 2A,B) mostly showed a linear trajectory without saturation or reset of the types shown in Figure 3 (left). The continuation of the accumulation without resets may be a result of the active tracking. We do not know precisely why this occurred but we speculate it was either because 1) the extra attention required to track decreased the chance of attentional distraction introducing resets (Nakayama & Holcombe, 2020) or 2) the temporal demands of the drawing the trace in real time caused the participants to miss the resets that did occur, or 3) the resolution of online tracing at these speeds is poor and participants may have just averaged their traces over any breaks. In addition, post-dictive inference may prevent resets from being perceived. To address the possibility that it was the active tracking that suppressed the resets, we switched to a delayed, offline recording of the perceived path and the participants’ trajectory now showed frequent resets.

We were interested, in particular, in whether the limitation of the accumulation would be set by space or time. Since both the true and the illusory position appear to be available in the visual system (the true position drives saccades whereas the illusory position drives perception, Lisi & Cavanagh (2015)), it is possible that resets occur once the distance between the real and perceived positions exceeds some limit. We can speculate that this spatial limit would be related to the positional uncertainty of the gabor. When the position information is reliable, the resets would occur with very little illusory deviation, keeping the perceived path close to the true path. When the positional uncertainty is high, in the periphery, with the gabor’s mean luminance matched to the background (Cavanagh & Tse, 2019; Gurnsey & Biard, 2012), the spatial offset could be quite large before exceeding the range of positional uncertainty around the gabor’s true position. Under this hypothesis, the occurrence of spontaneous resets will depend on the spatial offset from the true path which will be directly proportional to the internal speed (for a constant external motion). On the other hand, if spontaneous resets are the result of the temporal limitations of the integration process, resets would occur after a certain amount of time, independently of the internal speed and the spatial offset it creates. Investigating the time and location of resets in the recorded trajectories, allows us to distinguish between a spatial limit to the illusion and a temporal limit. This in turn will inform us of the processes underlying the perception of the position of moving objects.

To do so, we ask participants to re-trace the perceived path of a double-drift stimulus, immediately after viewing the stimulus, a method used by Nakayama & Holcombe (2020). This allows participants more time to carefully reproduce the path. We use the recorded trajectories to assess points where resets occurred, whether they were a discontinuity in orientation reflecting a saturation (hit the wall reset, Fig. 3) or a discontinuity in position reflecting a jump back to the real location. These reset points can then be used to determine if resets are time-limited or space-limited.

### Methods

#### Participants

For this experiment, 9 participants were recruited from the lab (6 female; ages 19 - 27, mean: 22.8). All participants reported themselves as right handed and had either normal or corrected-to-normal vision. Procedures were in accordance with the Declaration of Helsinki (2003) and were approved by York’s Human Participants Review Committee. All participants provided prior, written, informed consent.

#### Setup and Stimuli

The setup was the same as used in the online tracking experiment, except that we now also used a small keypad for responses (Fig. 1, right). Stimuli were identical to those used in the online tracking experiment, but now always started at the near end of the workspace and moved away from the participants. The double drift stimuli would use either 3 or 4 seconds to move 13.5 *cm*, corresponding to speeds of ∼4.5 and ∼3.75 *cm/s* or ∼4.7 and ∼3.5 *dva/s*, and had an internal drift of 2, 3 or 4 *cps*.

#### Procedure

Before the experiment started, the illusion and its resets were explained to particpants, and they were instructed to make sure to replicate any changes in direction of the perceived path as accurately as they could manage. In experiment 2, participants first peripherally observed a double-drift stimulus, while fixating to the left or right of the stimulus. Then they either reproduced the perceived path of the stimulus by retracing it on the tablet, or indicated the initial movement direction by changing the orientation of a line originating in the same position as the doube-drift stimulus (data not used, but corresponds to illusion strength determined from trajectories). More details are given below. This provided more time for the participants to reproduce any resets, i.e. without the need for real-time tracking. The re-tracing task was done in half of 8 blocks (the other half of the blocks are not used). Block types were alternated, and the order was counterbalanced across participants. Each block used all 6 combinations of two external speeds (corresponding to 3 or 4 *s* presentation time), and three internal speeds (2, 3 or 4 *cps*), 6 times, for a total of 36 trials per block. Each of the combinations of internal and external speed was presented 24 times in total, so that we had 144 trials for each participant, and 1296 trials in total.

In both kinds of trials, participants first had to move the stylus to the start position of the gabor, and then fixate a point to the left or right of the centre of the workspace. The double-drift gabor started at the near end of the display, and moved along the horizontal display away from the participant, always making a single, forward pass. After the gabor had disappeared, participants could respond in one of two ways. When re-tracing the perceived path with the stylus on the tablet, the drawn path would show up as a red line. Participants could “key in” their response by pressing ENTER, or start over by pressing ESCAPE. There was no time-limit for the response. Re-tracing the perceived paths should allow capturing spontaneous resets of the illusion, if there are any.

#### Analyses

The individual trajectories revealed inflections of direction that were potential spontaneous resets and we created a heuristic to localize them. First, we removed the first and last 4 mm of the trajectory, as well as any short segments on the trajectory at the start and end that went toward the participant (i.e. more than 180 degrees away from the true external motion), to remove any unintended jitters caused by starting and ending the movement. Then, we applied a bi-directional low-pass Butterworth filter with a cut-off frequency of 1.5 Hz. However, since participants draw at very different speeds, we resample them at 30 Hz using linear interpolation, giving 90 samples for the 3 *s* trajectories and 120 samples for the 4 *s* trajectories. After filtering, we detected the first inflection point in these summary trajectories using only the x coordinates (local maximum of x). This could put the reset point too late for saturation or “hit-the-wall” resets, so that we moved back to get the previous sample at 95% of the x coordinate at the inflection point. Then, we located the sample in the raw trajectory that was closest to this point. We then excluded a few reset points that were less than 5 mm, within the expected quadrant. Those that were very close to y<5 *mm* are unlikely resets because they either have extremely high illusion strength, or not enough room or time for actual accumulation of illusory drift. Those that were very close to the veridical path (x<5 *mm*) likely don’t have any illusory drift, and were probably random wiggles in an attempt to make a vertical tracing. Out of 1296 trials, the heuristic identified reset points in 827 (∼64%). This heuristic likely yielded some misses and some false alarms, and while changing the algorithm may improve performance, there is no ground truth to evaluate this. All raw trajectories and the detected reset points could be found on the OSF repository.

The X coordinates of the detected reset points give the deviation from the vertical path and are used as the spatial offset of the reset. We also need the time of the reset. The gabor moves at a uniform speed along the Y axis so distance along Y denotes the elapsed time for the physical path. However, the illusory path of the gabor does not have to stay in alignment with its physical location on the Y axis. If it did, its speed and path length would have to increase for larger illusion angles: for example, 41% faster and farther for a 45° illusion, 73% faster and farther for a 60° illusion. Observers have not reported these increases in speed or path length compared to the control with no internal motion. Moreover, in a recent unpublished study we directly measured the perceived path length of the illusion. The path length was always fixed proportion of the physical path (around 70%) for all illusion strengths, Gabor speeds and physical path lengths. This means that the perceived speed along the illusory path must be a fixed proportion of the physical speed as well and therefore, in the tracings, it is the distance along the traced path (the radius in our diagrams) that tells us the time since the trial began — not the distance along the physical path (the Y axis).

To facilitate modeling, the horizontal (illusory deviation) and vertical (straight ahead) coordinates of the reset points are kept isometric by giving them in centimeters. The distance from origin can then be converted to time as 1 *cm* = 4/13.5 *s* in the 4 *s* condition and 1 *cm* = 3/13.5 *s* in the 3 *s* condition. A probability density function is used to estimate each model’s likelihood and the variable that is used (X or T) is divided by its median value so that each of their probability densities are on a comparable scale.

Apart from reset points, we also extracted a measure of illusion strength. We took a trajectory sample at half the distance between the starting point and the detected reset point. We then used the angular difference between the gabor’s real trajectory and a straight line drawn through that point and the start of the trajectory as a measure of illusion strength.

#### Results

Four example trajectories and the location of the first resets determined by our heuristic are shown in Figure 4 (A-D). The x-axis here indicates illusory drift, since the gabor moved with a constant speed along the y-axis. Distance to origin can be converted to the time of reset by dividing by speed. On average, our heuristic detected resets in ∼64% of trials. For two participants, less than half the trials showed resets (9% and 22%). For the remaining seven participants the algorithm detected resets in 65%-99% of trials, with an average of 82%. Since participants experiencing a “hit-the-wall reset” (Fig. 3) as opposed to a “jump” reset, might not always display a detectable direction change in their paths, most of these percentages of trials with resets seems reasonable. However, the participant with only 9% of trials showing a reset was excluded for calculating illusion strength as they had no resets in several conditions. This means we analysed a total of 827 reset points (Fig. 4E).

**Figure 4:**
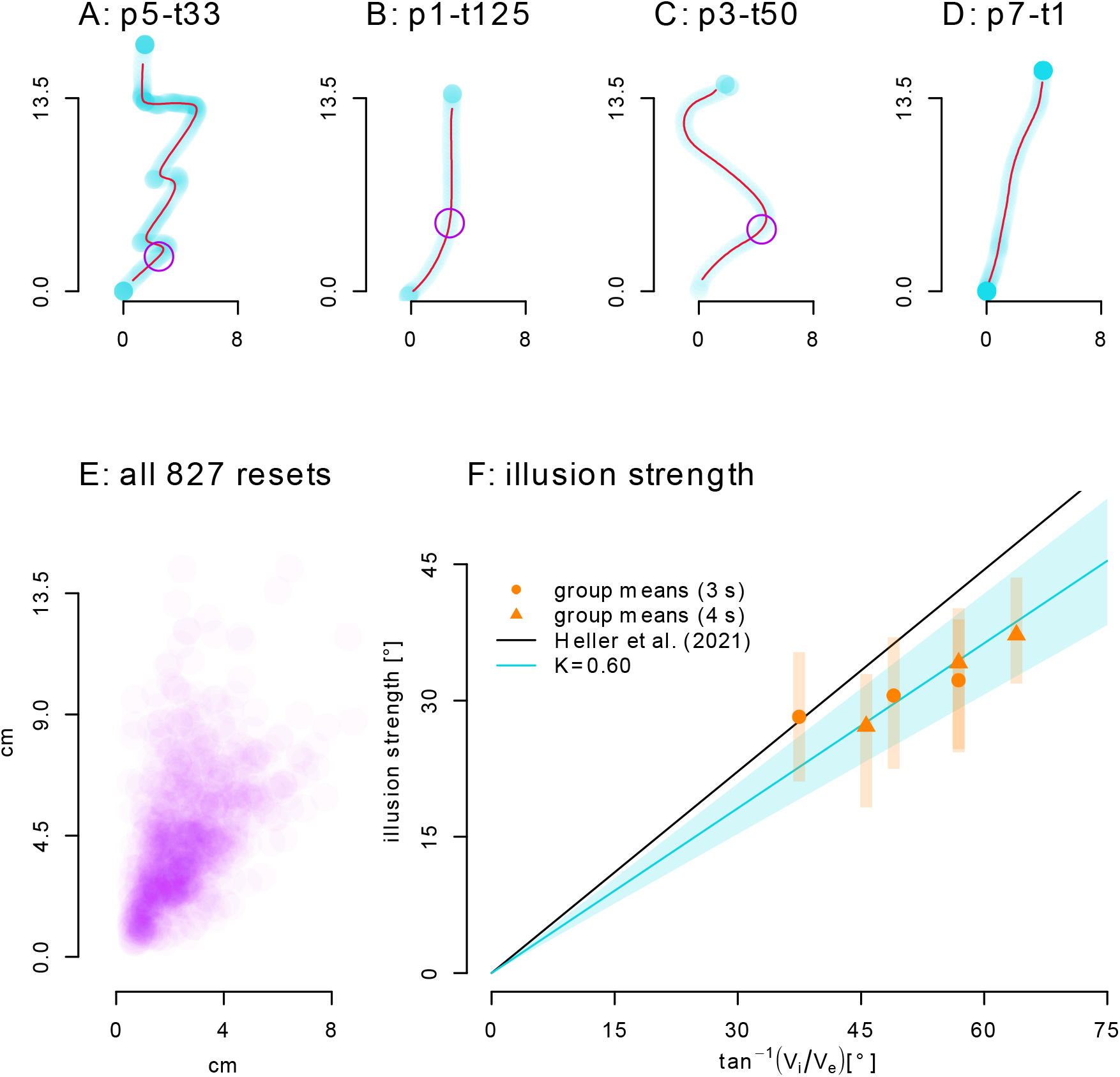
Re-tracing Data. **A-D:** Example trajectories and reset points. All trajectory samples in transparent light blue (denser samples result in darker color). Dark red lines are, pruned, interpolated and filtered trajectories (see Methods). Purple circles indicate reset points. **A:** Trajectory with several sudden resets. **B:** ‘Hit-the-wall’ reset. **C:** Slow return reset. **D:** Continued accumulation of the illusion without any reset. **E:** Distribution of all 827 detected reset points. **F:** Using the angle from the origin of the trace to a point along the trajectory at half the distance between origin and reset point as a measure of illusion strength, we can compare these data to other data. For this graph, we exclude one participant who produced very few resets. The illusion is weaker than in Experiment 1 (K=0.60, blue line) and its strength varies between participants (from K=0.40 to K =0.81), but it is nevertheless clearly there.

As before, we compare the strength of the illusion (Fig. 4F), with an earlier study (Heller et al., 2021). A line through the origin with *K*=0.60 predicts illusion strength fairly well in these data, which is weaker than the *K* =0.74 found by Heller et al. (2021). Illusion strength is also lower than in Experiment 1 with online tracking and it has a fair amount of between-subject variation in this task (blue shaded area is 95% confidence interval on K across participants, and orange lines are 95% confidence intervals across participants for illusion strength in each condition). While the illusion, as captured by retracing, is less strong, it is nevertheless there and depends on the relationship between internal and external speed as in previous studies.

#### Limits on the Double-Drift Illusion

Now we look at the distribution of reset points, and whether they can be explained by a temporal or spatial limit. In Figure 4E, we can see that the reset points vary more along the y-axis, than along the x-axis. This would be consistent with a spatial limit to the illusion resulting in resets. However, there is also considerable variance along the x-axis, so that fitting any vertical line through the data, (a spatial limit, the blue line in Fig 5A) provides a poor explanation of the pattern of resets.

**Figure 5:**
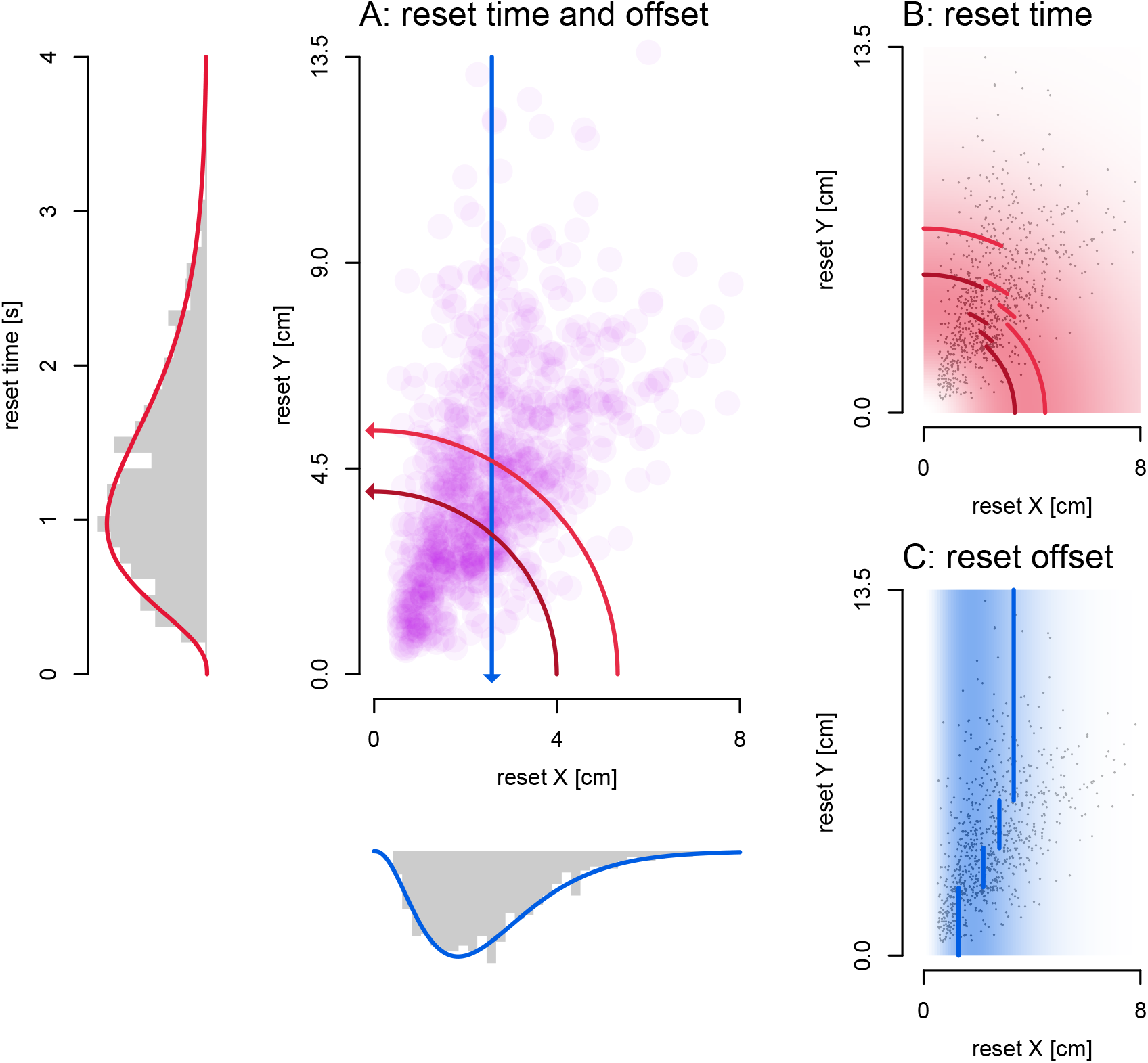
Reset models. **A:** Model limits relative to reset point locations. Reset points neither fall close to a spatial limit (blue line) or a temporal limit (dark red line: 4 s passes, light red line: 3 s passes). Marginal distributions show that both the time and spatial offset of reset points approximate a gamma distribution. **B**,**C:** Color illustrates the gamma probability distributions, where white is zero probability, and the darkest red or blue is the maximum probability density. **B:** Gamma distribution on time (AIC = 1215.12). Dark and light red lines: denote the median reset time for 4 s and 3 s passes in angular bins with an equal number of resets. **C:** Gamma distribution on spatial offset (AIC = 1269.18). Blue lines indicate median reset offsets in Y-coordinate bins with an equal number of resets.

We examined 4 simple models. If there is a spatial offset limit that triggers resets, the reset points should fall on, or be close to a line with a constant x coordinate (blue line in Fig. 5A). If there is a temporal limit, all resets should have the same or a similar reset time (distance from origin / speed, the red arcs in Fig. 5A). If we assume that these spatial or temporal limits do allow a little random variation from trial to trial, we can fit a normal distribution (with 2 parameters: mean and standard deviation) on both the x coordinates and the time points of resets. However, neither the x coordinates, nor the time points of resets were normally distributed (both *p*<.001, Kolmogorov-Smirnov tests of data against optimal normal distribution). Additionally, if resets are triggered by either a spatial or temporal limit the data would all fall relatively close to it, but this does not appear to be the case.

These first two models assumed that resets were triggered at fixed locations in space or time, but the next two propose that the resets are random processes that occur with some distribution across space or time. We used gamma distributions to capture these processes (these have 2 parameters: rate, the chance of an event per second; and shape, how many events are needed, in this case, until a reset is triggered) on the X coordinates and time points of resets as well (see Fig 5A, marginal distributions; Fig 5B,C). To compare across the four models, we then calculated the likelihoods of their predictions (based on the underlying distributions’ probability density functions) using all resets together and also within individual participants. From these likelihood values, we calculated an AIC to determine a relative probability for each model.

For the space and time limit with normal distributions (means in Fig 5A, blue: spatial, red: time), the time-based distribution fits better for the group data (normal space *AIC*=2,834.83, normal time *AIC*=1,677.98; *p*<.001) as well as all individual participants, and the same holds within gamma distribution models of reset models (gamma space *AIC*=1,269.18, gamma time *AIC*=1,215.12; *p*<.001). Furthermore, the gamma-based distribution on time of resets performed better than the normal distribution of reset time in the pooled data (*p*<.001), and this holds for most participants (6/9) as well. This means that time-based distributions explain our reset points better than models assuming a spatial offset, so that the time-based gamma distribution was the most successful of all 4.

While the marginal distribution plot in Fig 5A qualitatively appear an equally good fit to the data, the fit for the distribution on the time of resets maintains a better fit throughout the data set. We demonstrate this by dividing the data into segments, each with an equal number of resets, based on variables orthogonal to the fitted variable (Y-coordinates for the spatial offset of resets and angle for reset time). The median for each segment of data is shown in Fig 5B,C. We fit a gamma distribution to each segment, then use the data points outside the segment to get a log likelihood as a measure of generalization and sum them. We use this to calculate AICs and a relative likelihood (spatial: 6191.7, temporal: 3628.6, *p*<.001) which shows that a gamma distribution fit to reset times in one segment of the data generalizes better to the rest of the data, as compared to a gamma distribution fit to the spatial offset of resets.

## Discussion

We first tested whether manual tracking of a double-drift stimulus is susceptible to the illusion as reported by perceptual judgments, pointing (Lisi & Cavanagh, 2017), and memory saccades (Massendari et al., 2018; Ueda et al., 2018). The results of this online tracking experiment showed a close agreement between the angle of the manual tracking and the illusion strength expected based on the model from earlier perceptual measures (Heller et al., 2021). However, we were also interested in spontaneous resets and only ∼4% of trials in this first experiment had deviations that could have been a reset. In contrast, in the second experiment, 63.8% of trials showed a reset. There was a reset within 1 *s* in 22.3% of trials, and within 2 *s* in 52.2% of trials. We can use these results to determine the rate of spontaneous resets that would have been expected in the first experiment if it had the same reset dynamics as the second experiment. If each pass was independent and the perceived trajectories were equivalent in the two experiments, this would mean ∼98.5% of trials of Experiment 1 should have shown at least 1 spontaneous reset which is far larger than the ∼4% of trials we observed (*p*<.001, exact binomial test). We are not sure why so few spontaneous resets occurred in the first experiment. We first suggest two simple explanations for the difference: 1) Online tracking required relatively fast hand movements, and therefore left little time for drawing the details of any spontaneous resets. 2) The demands of the online tracking increased the attention to the double-drift stimulus and with less chance of attentional distraction, the likelihood of resets was reduced (Nakayama & Holcombe, 2020). However, this task, unlike previous tests of the double-drift stimulus, may also involve an interaction between action and perception (e.g. Beets et al., 2010). For example, the predicted sensory consequences of the hand movement may influence perception (“I will move my hand in this direction, so I expect to see motion in that direction as well”). Or vice versa, as in post-dictive inference (“my hand has moved in this direction so I must have seen the visual stimulus move in the same direction”). But this is a question for future research.

In our second experiment, we asked participants to draw the perceived path of the stimulus after observing a single pass. The orientation of the drawn traces reflect the strength of the illusion, and in a majority of trials (63.8%) there was a clearly indentifiable reset point, of either a “return” or a “hit-the-wall” type. We then set out to test if these spontaneous resets were triggered by a limiting distance of drift away from the physical location, or by a specific limit of time. Neither of these provided a viable explanation of the pattern of resets. Resets points are broadly distributed, suggesting that resets are not linked to the degree of illusory offset or to the time from stimulus onset, but instead may be caused by a more “random” process. Overall, both the spatial offsets and time of resets seem to follow a gamma distribution about equally well (Fig 5A, marginal distributions). However, average spatial offsets seems to be lower with lower Y coordinates, such that the gamma distribution on spatial offsets doesn’t work as well throughout the data set (Fig 5C). In contrast, a gamma distribution on reset time works equally well, irrespective of angle (Fig 5B), and this is reflected in the better fit of this model. Hence it seems most likely that reset points are randomly distributed over time, according to a gamma distribution.

Recently, it has been suggested that resets can be triggered by a distraction of attention (Nakayama & Holcombe, 2020). The participants in their experiment also retraced the perceived path and reported “return” resets. Nakayama and Holcombe (2020) claim that planned eye movements can trigger a “return” reset back to the physical path. In our experiment, we did not control for eye-movements, so we can’t directly investigate whether eye movements accompanied the “return” resets that participants reported. However, the distribution of resets throughout time are roughly in line with a mechanism based on random distractions of attention, which may involve eye-movement planning and execution. The rate of resets that we found here, ∼0.75 / *s*, is too infrequent for microsaccades (∼3 / *s*) so the distractions may be more significant than these small eye movements. On the other hand, gamma distributions (e.g. McGill & Gibbon, 1965; Wolfe et al., 2002) do model a random occurrence of events (rate parameter) where only some fraction of these trigger an action (shape parameter). In other words, the good fit of a gamma distribution may reveal some properties of the process underlying resets of the double-drift illusion. While the process giving rise to resets is likely memoryless, a process giving rise to a gamma distribution (with shape > 1) is not. Nevertheless, it does capture two ideas: 1) resets are triggered by random events (such as distractions of attention or microsaccades), and 2) only some of these actually lead to a reset. An alternative to the gamma distribution, the ex-Gaussian (e.g. Matzke & Wagenmakers, 2009; Guy et al., 2020), could be a more realistic alternative but does not fit the data as well. In any case, the process that actually leads to resets remains to be seen in future work.

The fit showed that reset times were well modeled by a broad gamma distribution, with resets occurring randomly in time (set by the rate and shape parameter) with an average rate of 1 reset every 1.3 *s* over the 3 to 4 *s* duration of the trials. However, we have to qualify our conclusions given the noisy nature of hand-drawn trajectories as a measure of the perceived trajectory and the reset points. To gauge the generality of our results, we compared them to those from a similar study that analyzed trajectory endpoints (pre-print: Liu et al., 2021) to determine the presence and size of spontaneous resets. In their study, the internal speed of the Gabor texture was adjusted across conditions and participants so that the illusion always appeared locally downward (vertical, ignoring any resets) compared to its actual 45° direction. This generated a highly predictable offset of path length /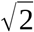 at the end of the trajectory if there were no spontaneous resets. Any measured offset less than the expected value (path length /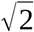) then revealed the presence and size of spontaneous resets. Liu et al. (2021) used this technique to measure spontaneous resets for 3 different durations and 2 path lengths (see Fig. 6A).

**Figure 6:**
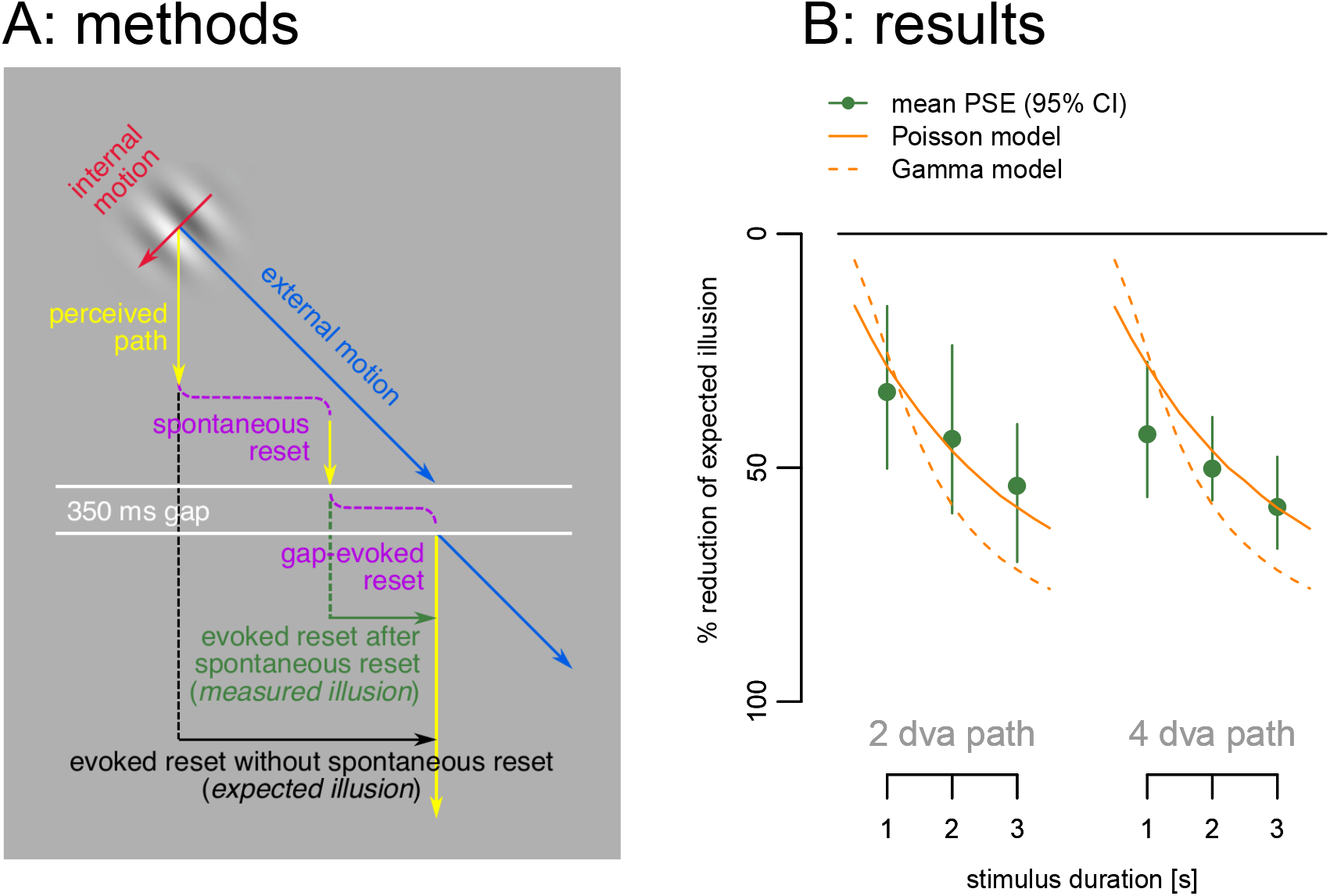
Spontaneous resets without hand trajectories from Liu et al. (2021), re-analyzed here. **A:** The Gabor’s internal motion was adjusted for each condition and participant so that the perceived path was always locally straight down (yellow arrows). The 350 ms temporal gap (white lines) then ensured a full reset (Lisi & Cavanagh, 2015) to the physical location when the Gabor continued on its path following the gap (blue arrow). Gaps were presented after 1, 2 or 3 seconds, and 2 or 4 dva real paths and the offset measured between the pre- and post-gap paths (vertical green line) indicated the amount of illusion left at the end of the pre-gap path (measured illusion), which can be contrasted with the expected illusion if there were no spontaneous resets (black line). **B:** Percent reduction of the illusion (difference between measured illusion and the expected illusion with no intervening spontaneous resets as a percentage of expected illusion) as a function of path length and duration. The size of the illusion decreased over time for a fixed path length and local illusion angle (green dots). These data are consistent with resets occurring randomly over time at ∼0.7 / s modeled as a Poisson process (continuous orange line), and largely with the gamma distribution fit to the reset times found here (dashed orange line).

The data from 10 participants showed that for a fixed path and illusion strength, longer durations produced smaller illusory offsets at the end (Fig 6B, green dots). This can be explained if spontaneous resets occurred randomly over time, so that longer durations would have more resets and a smaller final offset at the endpoint. Liu et al. (2021) modeled this with a Poisson process having a constant chance of reset of ∼0.7 / *s* (Fig 6B, orange line). In comparison, in our experiment, the gamma distribution fit to the data of explicit resets (rate=2.81, shape=3.74), gave a ∼0.75 / *s* chance of a reset, a remarkably similar estimate. We then used the parameters of our gamma distribution fit to our data to generate the PSEs that should be found in Liu et al.’s experiment with no additional fitting (Fig 6B, dashed orange line). We see that the two findings, based on very different approaches, are mostly in agreement with each other.

Kwon, Tadin, and Knill (2015) proposed a Bayesian object tracking model to explain the double-drift illusion and other phenomena. In their model, when the position signals are weak they have less weight and the predictions based on motion signals get more weight, moving the perceived position away from the true location. Their model predicts that after about 200 *ms* the motion prediction saturates and the perceived path then continues parallel to the physical path, as in the “hit the wall” resets we describe above (Kwon et al., 2015, Fig. 3). In contrast to their model, the resets in our re-tracing task show quite variable timing (Fig. 5A) and are better described by a fairly broad gamma distribution (Fig. 5B). This argues against a mechanism that is based on comparing the current location to the physical location and saturating or triggering a reset when the discrepancy is too large. In any case, the best fitting temporal limit of the reset points in our data was not 200 *ms* but 1.3 *s* and 99% of detected reset times are above 200 *ms* (see Fig. 5A). Moreover, 4/9 of our participants clearly reported “return” resets that the Kwon et al. (2015) model does not predict, and only 1/9 participants showed clear “hit-the-wall” resets. The “return” resets suggest that both the physical and perceived locations are available to the visual system and that when spontaneous resets occur, the perceived location returns toward the physical location. Our results suggest that the resets, whether saturation or return resets, happen randomly over time, rather than at a particular deviation in spatial offset or after a fixed duration. Future models of position perception for the double-drift stimulus need to include a realistic model of the locations and type of resets.

In summary, we find that manual tracking of the double-drift stimulus during its motion did show the expected illusion at the expected strength (experiment 1). Few if any resets were detected during online tracking, however, perhaps because of the additional attentional requirements or the short duration (2 *s*), although interactions with the simultaneous hand movements may contribute as well. In contrast, participants did report relatively frequent, spontaneous resets in the delayed, single-pass re-tracing of experiment 2. The resets were best explained by a gamma distribution on reset time. This finding suggests that spontaneous resets occur independently of the stimulus and how or where it is perceived. Instead, they may occur randomly over time, perhaps due to unrelated attentional distractions.

## Acknowledgements

The research was support by funding from NSERC Canada (PC and DYPH).

## References

Arnold, D., Thompson, M., & Johnston, A. (2007). Motion and position coding. Vision Research, 47(18), 2403–2410. https://doi.org/10.1016/j.visres.2007.04.025

Beets, I. A. M., ‘t Hart, B. M., Rösler, F., Henriques, D. Y. P., Einhäuser, W., & Fiehler, K. (2010). Online action-to-perception transfer: Only percept-dependent action affects perception. Vision Research, 50(24), 2633–2641. https://doi.org/10.1016/j.visres.2010.10.004

Benjamini, Y., & Hochberg, Y. (1995). Controlling the False Discovery Rate: A Practical and Powerful Approach to Multiple Testing. Journal of the Royal Statistical Society. Series B (Methodological), 57(1), 289–300. https://www.jstor.org/stable/2346101

Blohm, G., Missal, M., & Lefèvre, P. (2003). Smooth anticipatory eye movements alter the memorized position of flashed targets. Journal of Vision, 3(11), 761–770. https://doi.org/10.1167/3.11.10

Cai, R., & Schlag, J. (2001). Asynchronous feature binding and the flash-lag illusion. In Investigative Ophthalmology & Visual Science (Vol. 42).

Cavanagh, P., & Anstis, S. (2013). The flash grab effect. Vision Research, 91, 8–20. https://doi.org/10.1016/j.visres.2013.07.007

Cavanagh, P., & Tse, P. U. (2019). The vector combination underlying the double-drift illusion is based on motion in world coordinates: Evidence from smooth pursuit. Journal of Vision, 19(14), 2. https://doi.org/10.1167/19.14.2

Chung, S. T. L., Patel, S. S., Bedell, H. E., & Yilmaz, O. (2007). Spatial and temporal properties of the illusory motion-induced position shift for drifting stimuli. Vision Research, 47(2), 231–243. https://doi.org/10.1016/j.visres.2006.10.008

Cormack, L. (2019). Dynamics of Motion Induced Position Shifts Revealed by Continuous Tracking. Journal of Vision, 19(10), 294c–294c. https://doi.org/10.1167/19.10.294c

De Valois, R. L., & De Valois, K. K. (1991). Vernier acuity with stationary moving gabors. Vision Research, 31(9), 1618–1626. https://doi.org/10.1016/0042-6989(91)90138-U

Duhamel, J.-R., Colby, C. L., & Goldberg, M. E. (1992). The updating of the representation of visual space in parietal cortex by intended eye movements. Science, 255(5040), 90–92. https://doi.org/10.1126/science.1553535

Eagleman, D. M., & Sejnowski, T. J. (2000). Motion Integration and Postdiction in Visual Awareness. Science, 287(5460), 2036–2038. https://doi.org/10.1126/science.287.5460.2036

Gamble, C. M., & Song, J.-H. (2017). Dynamic modulation of illusory and physical target size on separate and coordinated eye and hand movements. Journal of Vision, 17(3), 23. https://doi.org/10.1167/17.3.23

Gurnsey, R., & Biard, M. (2012). Eccentricity dependence of the curveball illusion. Canadian Journal of Experimental Psychology, 66(2), 144–152. https://doi.org/10.1037/a0026989

Guy, N., Lancry-Dayan, O. C., & Pertzov, Y. (2020). Not all fixations are created equal: The benefits of using ex-Gaussian modeling of fixation durations. Journal of Vision, 20(10), 9. https://doi.org/10.1167/jov.20.10.9

Heller, N. H., Patel, N., Faustin, V. M., Cavanagh, P., & Tse, P. U. (2021). Effects of internal and external velocity on the perceived direction of the double-drift illusion. Journal of Vision, 21(8). https://doi.org/10.1167/jov.21.8.2

Hogendoorn, H. (2020). Motion Extrapolation in Visual Processing: Lessons from 25 Years of Flash-Lag Debate. Journal of Neuroscience, 40(30), 5698–5705. https://doi.org/10.1523/JNEUROSCI.0275-20.2020

Hui, J., Wang, Y., Zhang, P., Tse, P. U., & Cavanagh, P. (2020). Apparent Motion Is Computed in Perceptual Coordinates. I-Perception, 11(4), 2041669520933309. https://doi.org/10.1177/2041669520933309

Johnston, A., & Scarfe, P. (2013). The Role of the Harmonic Vector Average in Motion Integration. Frontiers in Computational Neuroscience, 7, 146. https://doi.org/10.3389/fncom.2013.00146

Knol, H., Huys, R., Sarrazin, J.-C., Spiegler, A., & Jirsa, V. K. (2017). Ebbinghaus figures that deceive the eye do not necessarily deceive the hand. Scientific Reports, 7(1), 3111. https://doi.org/10.1038/s41598-017-02925-4

Kwon, O. S., Tadin, D., & Knill, D. C. (2015). Unifying account of visual motion and position perception. Proceedings of the National Academy of Sciences of the United States of America, 112(26), 8142–8147. https://doi.org/10.1073/pnas.1500361112

Lisi, M., & Cavanagh, P. (2015). Dissociation between the perceptual and saccadic localization of moving objects. Current Biology, 25(19), 2535–2540. https://doi.org/10.1016/j.cub.2015.08.021

Lisi, M., & Cavanagh, P. (2017). Differential spatial representations guide eye and hand movements. Journal of Vision, 17(12). https://doi.org/10.1167/17.2.12

Liu, S., Tse, P. U., & Cavanagh, P. (2021). The perceived position of a moving object is reset by temporal, not spatial limits. bioRxiv, 472615. https://doi.org/10.1101/2021.12.14.472615

Massendari, D., Lisi, M., Collins, T., & Cavanagh, P. (2018). Memory-guided saccades show effect of a perceptual illusion whereas visually guided saccades do not. Journal of Neurophysiology, 119(1), 62–72. https://doi.org/10.1152/jn.00229.2017

Matzke, D., & Wagenmakers, E.-J. (2009). Psychological interpretation of the ex-Gaussian and shifted Wald parameters: A diffusion model analysis. Psychonomic Bulletin & Review, 16(5), 798–817. https://doi.org/10.3758/PBR.16.5.798

McGill, W. J., & Gibbon, J. (1965). The general-gamma distribution and reaction times. Journal of Mathematical Psychology, 2(1), 1–18. https://doi.org/10.1016/0022-2496(65)90014-3

Nakayama, R., & Holcombe, A. O. (2020). Attention updates the perceived position of moving objects. Journal of Vision, 20(21). https://doi.org/10.1167/jov.20.4.21

Nijhawan, R. (1994). Motion extrapolation in catching. Nature, 370(6487), 256–257. https://doi.org/10.1038/370256b0

Patricio Décima, A., Fernando Barraza, J., & López-Moliner, J. (2022). The perceptual dynamics of the contrast induced speed bias. Vision Research, 191, 107966. https://doi.org/10.1016/j.visres.2021.107966

Peirce, J. W., Gray, J. R., Simpson, S., MacAskill, M. R., Höchenberger, R., Sogo, H., Kastman, E., & Lindeløv, J. (2019). PsychoPy2: Experiments in behavior made easy. Behavior Research Methods, 51(1), 195–203. https://doi.org/10.3758/s13428-018-01193-y

R Core Team. (2019). R: A language and environment for statistical computing. R Foundation for Statistical Computing. https://www.R-project.org/

Shapiro, A., Lu, Z.-L., Huang, C.-B., Knight, E., & Ennis, R. (2010). Transitions between central and peripheral vision create spatial/temporal distortions: A hypothesis concerning the perceived break of the curveball. PLoS ONE, 5(10), e13296. https://doi.org/10.1371/journal.pone.0013296

Soechting, J. F., Engel, K. C., & Flanders, M. (2001). The Duncker illusion and eye-hand coordination. Journal of Neurophysiology, 85(2), 843–854. https://doi.org/10.1152/jn.2001.85.2.843

Tse, P. U., & Hsieh, P. J. J. (2006). The infinite regress illusion reveals faulty integration of local and global motion signals. Vision Research, 46(22), 3881–3885. https://doi.org/10.1016/j.visres.2006.06.010

Ueda, H., Abekawa, N., & Gomi, H. (2018). The faster you decide, the more accurate localization is possible: Position representation of “curveball illusion” in perception and eye movements. PLOS ONE, 13(8), e0201610. https://doi.org/10.1371/journal.pone.0201610

Wolfe, J. M., Torralba, A., & Horowitz, T. S. (2002). Remodeling visual search: How gamma distributions can bring those boring old RTs to life. Journal of Vision, 2(7), 735. https://doi.org/10.1167/2.7.735

